# A new model on DNA structure and thermal denaturation

**DOI:** 10.1101/386524

**Authors:** LUO Liaofu, YANG Guochen

**Affiliations:** School of Life Science and Technology, Inner Mongolia University of Science and Technology, Baotou, China; School of Physical Science and Technology, Inner Mongolia University, Hohhot, 010021 China; Hebei University of Technology, Tianjin, 300401 China

## Abstract

A quantum model on DNA structure is proposed. By introducing the self-consistent harmonic potential in the X-Y plane vertical to helix axis (z-direction) and the periodic potential along the z-axis we obtain the wave function for the single nucleotide and the many-nucleotide system. The helix distribution of bases is deduced from the solution of wave function under the self-consistent potential. The variation of DNA structure (polymorphism) is related to the periodicity of the potential in Z-axis, the quantum state occurring in harmonic potential and the interaction between helix strands. As Watson-Crick (W-C) interaction is introduced between double helices, the quasi-particle transformation is utilized to solve the interacting many-body problem for DNA. It is proved that the phase-transition (thermal denaturation) temperature is related to the frequency ω of harmonic potential. Through comparison with experimental data a simple relation 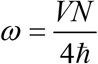 (*N* means number of base pairs and *V* the W-C coupling) is deduced. For a DNA sequence of 1000 bp ω is predicted about (0.9-1.2)×10^17^/sec. Such a high frequency is necessary for nucleotides of each strand located on a narrow tube. The large temperature fluctuation experimentally observed during DNA thermal denaturation is interpreted by the collective motion of nucleotides.

## 1 Introduction

Since the birth of molecular biology the quantum biochemistry has developed to treat the electronic motion in biological macro-molecules. However, the quantum dynamics was commonly ignored in studying problems such as biomolecular function and conformational change. Only in the latest years the quantum biology re-attracts again the attention of scientists. New progresses have achieved that include the non-trivial quantum effects in photosynthetic light harvesting [1,2], in avian magnetoreception [3] and in conformational transition of proteins[4,5]. However, to our knowledge, the quantum theory of DNA structure as an important part of quantum biology has not been found in literature. The motivation of the present studies is to filling up the gap and to constructing a quantum model of the DNA structure.

Life and non-life should have a unifying microscopic picture. The cornerstone of life -- atoms and molecules -- should obey unifying quantum rules. Although the QM/MM (quantum mechanics / molecular mechanics) multi-scale algorithm has achieved great successes in simulating the biological macro-molecule [6], the classical molecular mechanics (MM) is only an approximation in discussing the problems such as DNA structure. More rigorous studies based on quantum mechanics are necessary.

The miracle of heredity stability can only be explained by quantum mechanics. The point was indicated firstly by Schrodinger [7]. Since the energy level spacing of molecules participating DNA replication is larger than the thermal and other disturbances, the replication process will be stable. Nowadays, the epigenetic inheritance attracts the attentions of biologists [8]. Epigenetic inheritance describes different phenotypic consequences to be inherited without any change in DNA sequence. The stability of the epigenetic inheritance that is related to the existence of some self-perpetuating structure in an individual can still be explained following above demonstrations. Recently, the role of quantum decoherence was recognized by physicists and it is argued that the decoherence possibly makes the quantum picture ceasing to be effective for some macromolecular system. However, the nucleic acid molecule is a topologically ordered state with complicated non-local quantum entanglement. The non-locality means that the quantum entanglement is distributed among many different components of the molecule. As a result, the pattern of quantum entanglements cannot be destroyed by local perturbations. This significantly reduces the effect of decoherence and makes the quantum theory still effective in treating the genetic stability problems [9].

On the other hand, the study on the polymorphism of DNA structure has made great progresses in recent years. As is well known, even in DNA molecules of double helix, there are many structural types. Apart from double helix, it has been observed that the single-stranded DNA is generated for replication in phage, the triple helix and quadruple helix with couplings different from Watson-Crick types are also observed in experiments[10]. More interestingly, the transition of DNA conformational polymorphism was experimentally studied and detected on carbon nanotubes [11]. The polymorphism of DNA structure requires a unifying theory to interpret the mechanism for its formation. We suppose that the theory should be a quantum one, e.g. the nucleic acid had better be looked as a quantum complex system composed of many nucleotides.

The materials are organized as follows. In section 2 a model of nucleotide self-consistent-field is proposed and the solution of nucleotides in the field is obtained. Then in section 3 the interaction between double helices is introduced. The second-quantization method and the quasi-particle transformation are used to solve the many-body interaction problem. In the subsequent section 4 the thermal denaturation of DNA double-helix is studied and it is found that the transition temperature is related to the self-consistent field frequency. The observed temperature fluctuation during denaturation is explained. In section 5 the polymorphism of DNA structure is discussed from the solution of nucleotides in the self-consistent field. Finally in the last section, main results are summarized and several points related to the development of the model are discussed briefly.

## 2 Model: Nucleotide-self-consistent-field (NSCF) and the solution of nucleotides in NSCF

Nucleic acid is a system of multi-nucleotides. Although the quantum mechanical calculation for oligo-atom reactions was worked out and the rate constant was deduced in recent years [12], the generalization of the quantum calculation to the multi-atom system is still a difficult task. As a primary step to solve the problem we shall use self-consistent-field method. We model the single chain nucleotides as the independent fermions in a self-consistent-field of cylinder shape [13] The self-consistent-field can be deduced in principle through Hartree-Fock approximation if the interactions between these particles are known. However, we shall assume the self-consistent-field as a 2D harmonic potential or its shape- invariance set in XY plane and a periodic potential in Z-direction. That is, the single particle Hamiltonian written in cylinder coordinate is assumed as

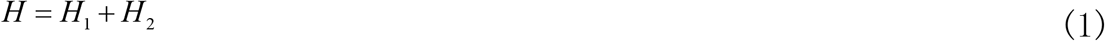

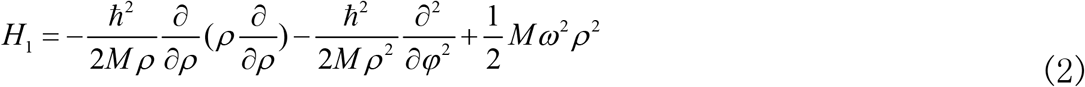

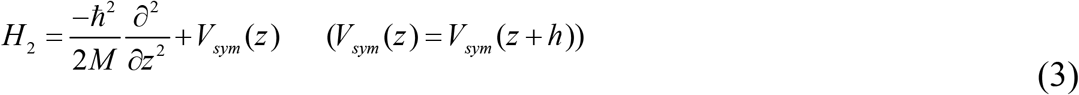

where *V_sym_* is periodic potential with period h.

The solution for Schrodinger Equation

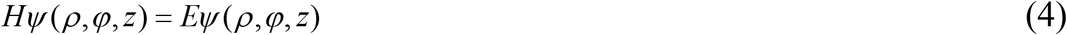

is

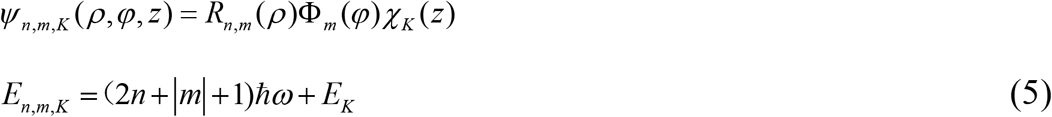

with *n*=0,1,2,…; *m*=0, ±1,±2,…. (Appendix 1) [14]. R_n,m_ (ρ) is related to a confluent hyper-geometry function that gives coordinate constraints on the wave function of 2D oscillator and

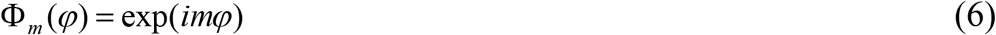

One may use another set of quantum number n_x_ and n_y_ to describe the state. n_x_ is the quantum number of x-direction oscillator, n_y_ is the quantum number of y-direction oscillator and one has n_x_ + n_y_ = 2n + |*m*|. The ground state of helix has *n*=0, *m* = 0 or n_x_=n_y_=0; the first excited state has two possibilities, *n*=0, *m* = ±1 or n_x_ =1, n_y_ = 0 and n_x_ =0, n_y_ =1; the second excited state has three possibilities, (*n*=1, *m* = 0), (*n*=0, *m* = ±2); etc. When *m* ≠ 0 it describes a helix state, when m=0 the state collapses to a line. The single chain is generally in the ground state of 2D harmonic SCF, while each strand of DNA is in the first excited state of 2D harmonic SCF. That is, as the Watson-Crick coupling is switched on, the nucleotides in SCF generally jump from ground state to the first excited state.

Following Bloch theorem we have

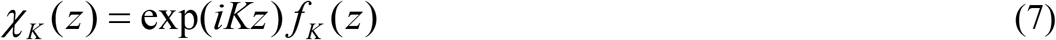

The subscript *K* is the Bloch wave number in OZ direction and *f_K_* (*z*) is a periodic function of z, *f_k_* (*z*)=*f_k_* (*z*+h). The energy *E_k_* describes the energy level in conducting bands and can be deduced from the periodic potential *V_sym_*(*z*).

Considering

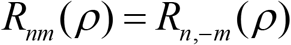

and

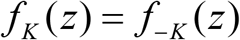

we obtain stationary-wave solution

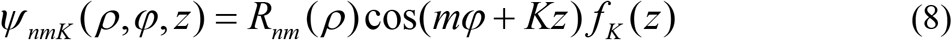

with the same eigenvalue given by (5).

In Equation (8) *f_K_*(*Z*) is a periodic function and the 2-dimenional wave function cos(*mφ*+*K_z_*) describes the helix distribution of bases in *φ-z* subspace with step of helix 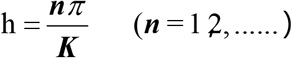. Moreover, the solution is invariant under the transformation

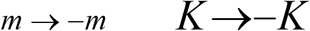

Suppose the system of *N* nucleotides is located in a cylinder of height (length) *L*. The Bloch wave-number is given by 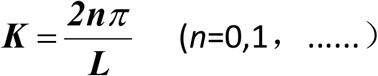. However for particles moving in a periodicity field the energy levels constitute several sets of conducting bands intermixed with forbidden bands. The boundary between a conducting band and its adjacent forbidden band (wave number denoted as K_b_) is determined by the condition [15].

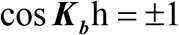

or *K*_b_h =*nπ* (*n*=l, 2,……). Only those K’s in conducting band are permitted. The observed step of helix in experiments is the average of 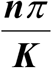 in some given conducting bands that is related to the breakdown of translational symmetry of the potential *V*_sym_(*z*).

### Remarks on second quantization and Wannier representation

For the further applications here we give two remarks. The motion of *N* nucleotides in a single strand can be regarded as a quantum many-body problem written directly by use of second quantization method. Define annihilation and production operator of the state labeled by quantum number *n,m,K* as

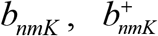

and assume they satisfy

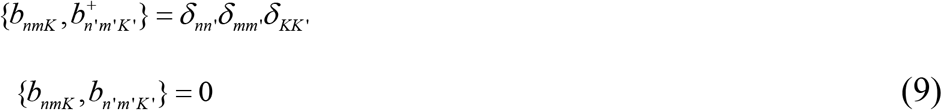

The many-body Hamiltonian of the system is

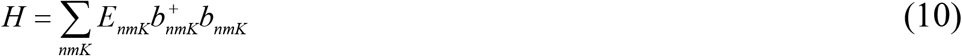

The annihilation and production operators satisfy Heisenberg’s motion equation.

Assume the nucleotides move in a periodic self-consistent field and the helix is composed of many primitive cells characterized by the lattice position **z**_*l*_. The *l*’s are taken from 1 to N (N cells) or from 1 to L/h (L/h cells). To be definite, we take (*l*=1,…,N) in the following. Define the Fourier’s transformation of Bloch wave function χκ(z)

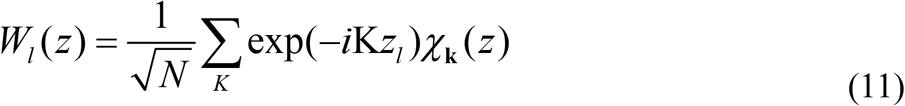

as the Wannier wave function in the periodic potential [16]. Under the tight binding approximation one can deduce a relation between the Bloch wave function in periodic field and the localized bound state wave function *φ_i_*(with energy level *ε_i_*,) [17],

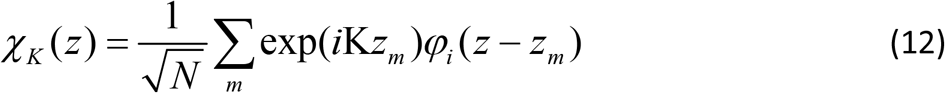

Eq (12) is exactly in the form of the inverse transformation of (11). So, the Wannier wave function *W_l_*(*z*) is essentially the locally bound state wave function *φ_i_*

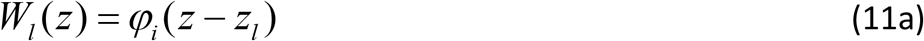

By use of (7) and (12) one has

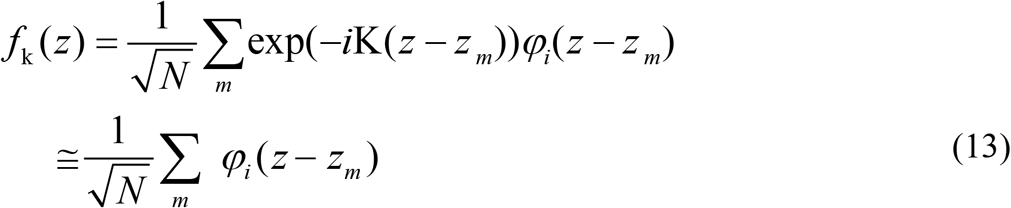

due to *φ_i_* not vanishing only as z near z_m_. It gives an approximate expression for Bloch wave function *f_K_*.

Generally speaking, the N-degenerate bound energy levels *ε_i_*, of the N-particle system form a band in the periodic field. However, the translational symmetry (periodicity) of *V*_sym_(*z*) is generally broken. It was proved that the small breakdown of the symmetry would cause the strong localization of the solution [18]. This is the so-called Anderson localization. Therefore, the tight binding model is really a good approximation in the present problem. Under local approximation the energy band of N-particle system degenerates to N energy levels with the same value *ε _i_*. Above deduction shows that the Wannier wave function is the eigen-function of H_2_ (Eq(3)) under the tight binding approximation. Based on Wannier wave function, the second quantization of the many-body problem in local representation can be deduced immediately. The annihilation and production operators in Wannier representation are denoted by *b_nml_*, 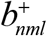 and they satisfy the same relation as Eq (9).

## 3 Model: Base pair interaction, double helix and quasi-particle

The formation of double helix must depend on the interaction between bases. Although each strand is localized in a small radius around an axis, the double helix extends to a larger spatial range due to Watson-Crick coupling. To understand the structure and dynamics of DNA one should study base pair interaction first.

By use of the annihilation and production operators in a many-body system the two-body interaction potential of the system can generally be expressed by

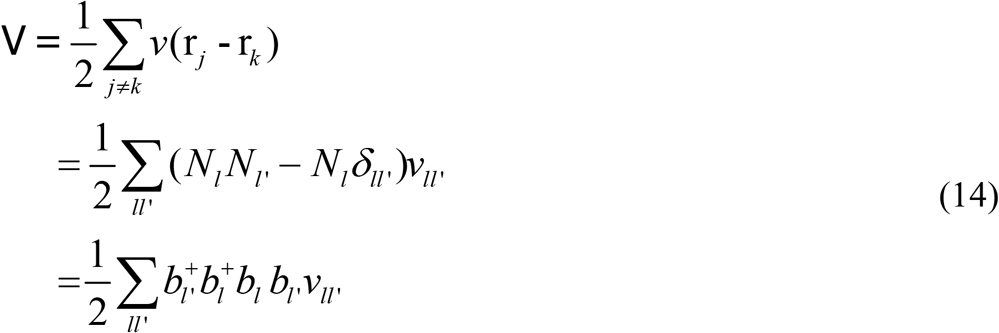

where *b_i_* and 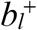 are the annihilation and production operator of quantum state *φ_l_* respectively and *N_l_*=*b_l_*^+^*b_l_* the particle number operator of the state *φl*. The 4-operator term in the last line of Eq (14) gives the second-quantization formalism for a general two-body interaction *ν*(**r_j_-r_k_**).

Set the annihilation and production operators on two chains in Wannier representation denoted by

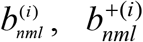

(the superscript *i*=1,2 means two chains respectively) and they satisfy the similar relation as Eq (9) in Bloch representation. The total Hamiltonian of the double helix system is given by

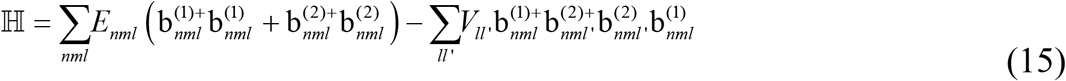

The second term in Eq (15) comes from Eq (14) which describes the Watson-Crick interaction between two chains and it can be rewritten as

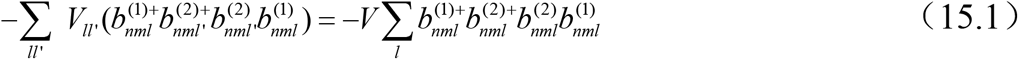

where only the interaction between corresponding bases (*l*=*l*’) on two chains is assumed and for simplicity the coupling is assumed independent of position *l*. Considering nucleotides on two chains are always in the same self-consistent field the same labels (*nm*) under all operators *b* and *b*+ will be omitted in the following. That is, *b^(i)^_nmi_* (*b^(i)+^_nmi_*) is abbreviated as 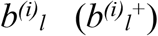.

As the W-C couplings switch on between corresponding bases of two strands the nucleotides of DNA are no longer in the quantum ground state *n*=0, *m*=0 but they jump from the ground state to the first excited state *n*=0, *m*= ± 1. The corresponding energy level of Wannier state is expressed by

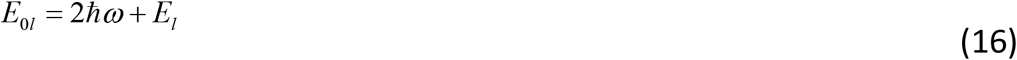

as seen from Eq (5) where *E_l_* is the energy of the locally bound state φ_l_ at z_i_. The reduced Hamiltonian of the double helix system is

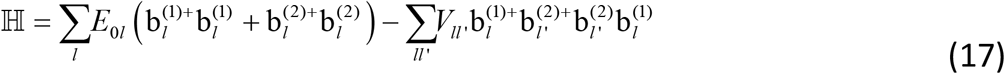

Due to the interaction term *V_ll’_* the Hamiltonian Eq(17) is not diagonal. However, the Hamiltonian can be diagonalized through a quasi-particle transformation as used in BCS theory of superconductivity [19]. Namely, we introduce the annihilation and production operator of quasi-particle

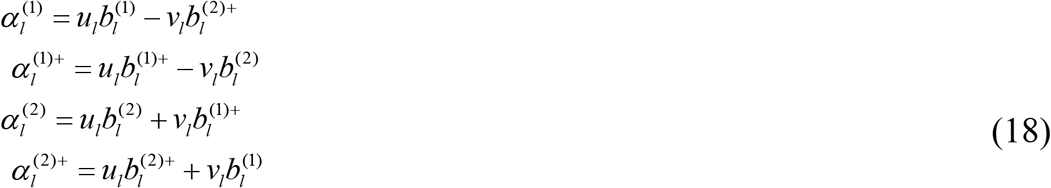

*u_l_* and *v_l_* are supposed to be real and satisfy

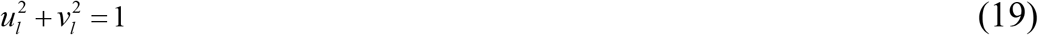

Easily prove 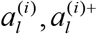 etc satisfy the Fermi type relation as Eq (9). So, 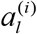 is the annihilation operator of quasi-particle and 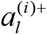 is the production operator of quasi-particle.

Set

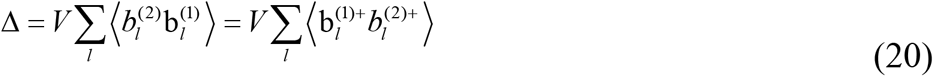

The reduced Hamiltonian of the helix system Eq (17) can be approximated as

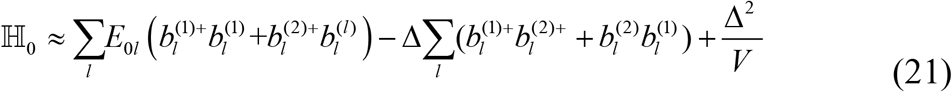

By transformation Eq (18) and under the choice of

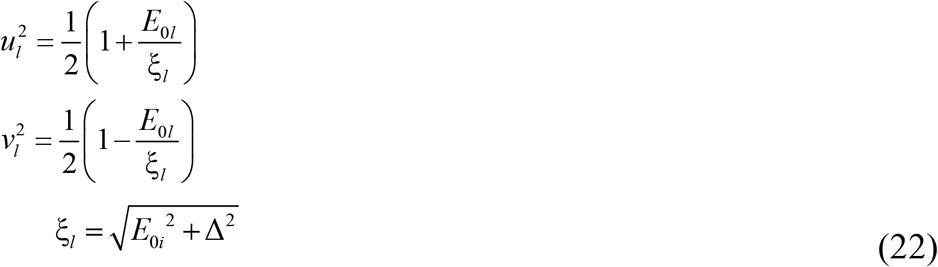

we obtain the Hamiltonian diagonalized relative to the quasi-particle operator

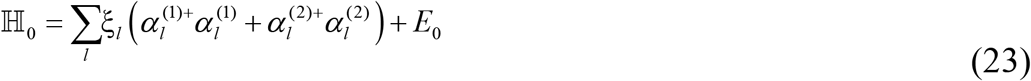

where

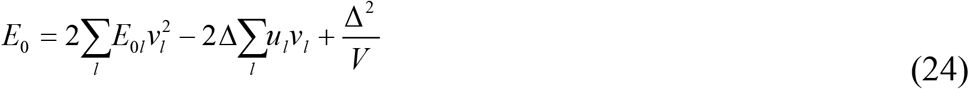

## 4 Thermal denaturation of DNA double helix

The phase transition of DNA occurs at the temperature *T_c_* where the gap Δ = 0. One should calculate the energy gap at first. At given temperature the thermal average of the gap, Eq (20), is expressed as

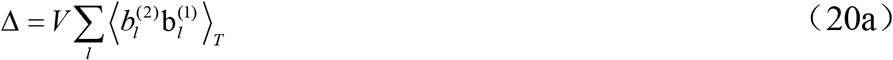

The inverse transformation of Eq (18) is

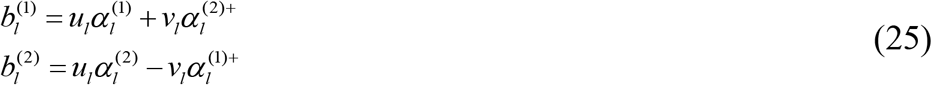

Inserting (25) into (20a) and neglecting two or more quasi-particle state we obtain

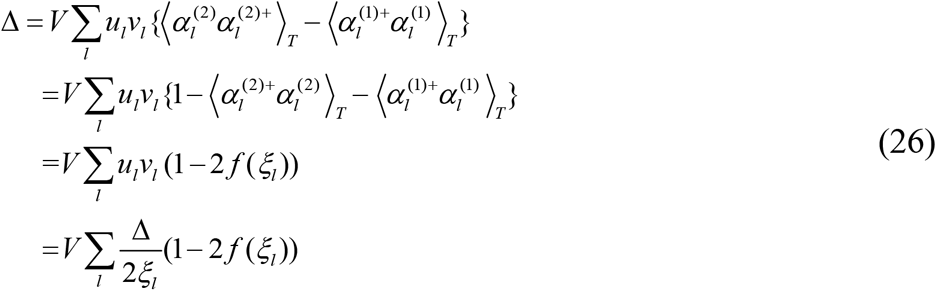

Here *f*(ξ_l_) means the distribution of ideal Fermi gas. In high-temperature and low density approximation, namely in case of the thermal wavelength much smaller than the average interparticle separation, the quantum effect can be neglected and the distribution *f*(ξ_l_) can be expressed as [20]

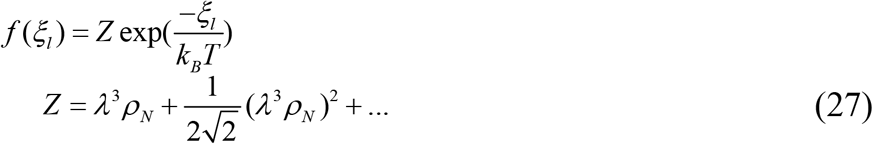

where *ρ_n_* denotes the number density of the particles and λ the thermal wavelength

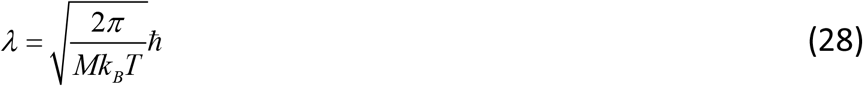

In experiments DNA undergoes a sharp transition at a definite melting point *T_c_* around 90°C [21]. The temperature Tc is high enough so that the condition λ^3^_ρN_<<1 is always satisfied in the problem. By using (26)(27) at *T*=*T*_c_ and taking Z<<1 into account we obtain

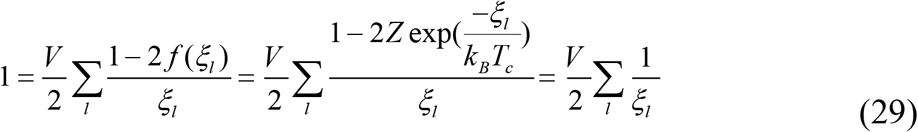

where *ξ_l_* = *E*_0*l*_ should be taken. It leads to

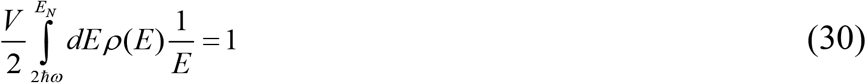

where the state density is given by (Appendix 2)

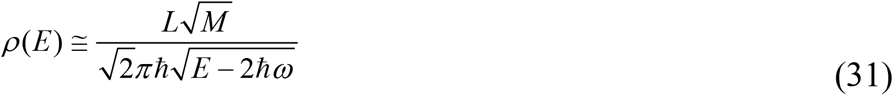

The energy integral in Eq (30) is completed in the harmonic-oscillator *n*=0, *m*= ± 1 band, namely, from the lower edge 2*ħω* to the upper edge *E_N_* approximately determined by

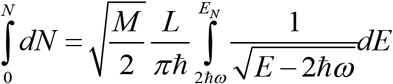

It gives

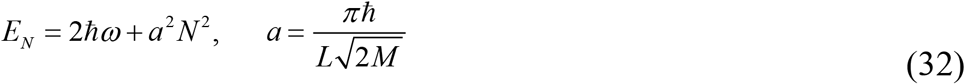

Inserting (31) and (32) into (30) one obtains

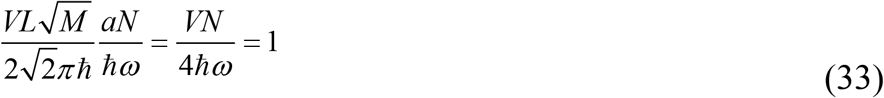

As an typical example, taking *N* = 1000, *V* = (0.35-0.48)×10^−12^*erg* corresponding to AT and GC coupling respectively we obtain

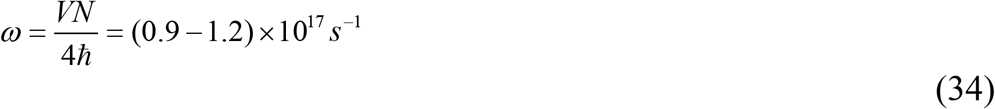

Above result holds as long as *Tc* is not-too-low so that Z<<1. It means the existence of phase transition at not-too-low temperature provides a clue to determine the frequency parameter of NSCF. The high frequency ω of self-consistent field is consistent with the idea that each strand of DNA is localized in a small radius that is comparable with the dimension of nucleotide 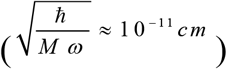.

The above discussion shows that the molecular mechanism of DNA thermal fluctuation is related to the collective motion of nucleotides in self-consistent field. Therefore, the denaturation temperature is dependent on the frequency of the field. The point can be further clarified by observing the fluctuation of denaturation temperature *Tc*. In fact, instead of Eq (33), more rigorous calculation gives

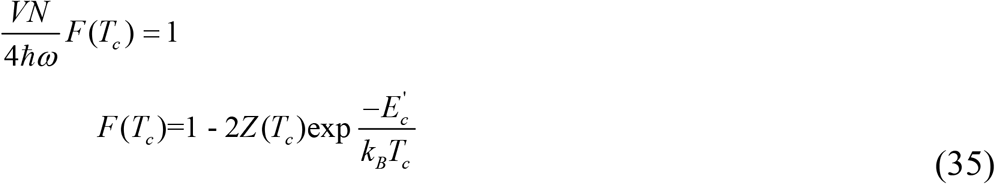

where *E’_c_* is an intermediate-value of *E* in energy integral from 2*ħω* to *E_N_* and (1-2Zexp(-*E*/*K_B_T_c_*) under the integral has been approximated by (1-2Zexp(-*E_c_’*/*K_B_T_c_*) and taken out of the integral. Both *N* and *V* have fluctuations. It leads to the fluctuation of *Tc*,

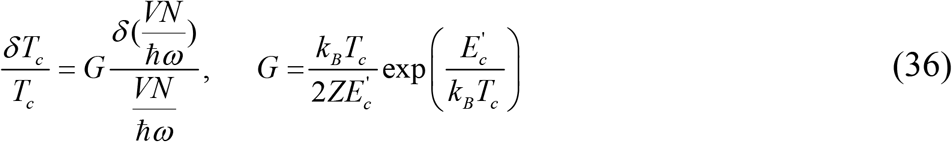

where *G*>>1 due to *E’_c_*> 2*ħω* >>*k_B_T_c_*. Therefore, the small fluctuation of N and V possibly causes the large fluctuation in temperature during DNA thermal denaturation This has been observed in experiments [22]. However, most part of fluctuations in N and V may have been canceled by the fluctuation of frequency ω. That is, the self-consistent field frequency ω is automatically adjusted to the fluctuated *N* and *V*. The observed temperature fluctuations are only the remainder of the fluctuation effect of N and V.

## 5 Polymorphism of DNA structure

There are three approaches to the variation of DNA structure. The first approach is due to the change of the quantum number (*n,m*) of the single chain in the two-dimensional harmonic self-consistent field. The quantum number *m*=0 describes the non-helix state, while the positive *m* and negative *m* describe right- and left-handed helix respectively. The quantum number *n* describes the nucleotide distribution in radial direction. Since ^2n + |*m*| = *n_x_* + *n_y_*^ where *n_x_* (*n_y_*) the quantum number of x-(y-) directional linear oscillator, the increase of *n* corresponds to the quantum state of particles distributed in larger x and/or in larger y-direction. These structural variation requires the large energy change which is in the order of ^*ħω*^.

The second approach to the variation of DNA structure is caused by the change of the symmetry existing in the Hamiltonian H_2_ (Eq (3)). For a single chain of B-DNA one helix turn (one step) contains 10 nucleotides, for A-DNA 11 nucleotides, for C-DNA 9.33 nucleotides, and for left-handed Z-DNA 12 nucleotides, etc. In solution the number of base pairs contained in one step of DNA further changes, for example, from 10.3 to 10.6 for B-DNA [10]. All above polymorphism of DNA structure can be understood by the periodicity of V_sym_(z) and the positive/negative directivity in φ-z subspace. In living cell DNA is winding on the histone octamer. The axis of DNA is a curve and in this case the tangent of the curve defines the z-direction. Suppose there are N nucleotides located in a tube-like region of length L and the self-consistent potential V_sym_(z) has symmetry V_sym_(z)=V_sym_(z+h). The step h of helix is determined by the interaction among N nucleotides and the interaction between nucleotides and environment. For a definite h the number of nucleotides in one step is changed as L stretched or contracted. It can be an integer or a fraction. Simultaneously, accompanying z increasing the angle φ may increases or decreases. These lead to polymorphism of DNA.

The third approach to the variation of DNA structure is due to the interaction between base pair in adjacent chains. The commonest structure is double strands paired through Watson-Crick coupling. In triple helix DNA the third strand is coupled with double helix through Hoogsteen or reversed Hoogsteen pairing. The quartet helix DNA is formed of four-stranded complex by guanine-rich motifs in DNA[10]. The theory of DNA thermal denaturation stated in previous section can easily be generalized and applied to the triple and quartet helix DNA.

## 6 Discussions

### 6.1 Main results

The present study is developed based on two basic assumptions: 1, The nucleotides in a single chain of DNA are independent fermions moving in a 2D harmonic self-consistent potential in the transverse plane and an 1D longitudinal periodical self-consistent field parallel to the helix axis. 2, The Watson-Crick interaction between double chains is described by the two-body interaction potential through the annihilation and production operators of nucleotides in the self-consistent field.

Based on above assumptions two main results are obtained. The first is: by introducing the self-consistent harmonic potential and periodic potential, the double helix structure is deduced and the polymorphism of DNA structure is explained or predicted. The second is: by introducing quasi particles in the coupled double helix the problem of DNA thermal denaturation is solved. The denaturation is a kind of collective motion of nucleotides, whose temperature is related to the frequency ω of harmonic potential. A simple relation 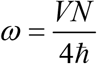 is obtained. We expect the relation will be further tested in future experiments.

### 6.2 Generalization from harmonic potential to self-consistent field of same shape

Set

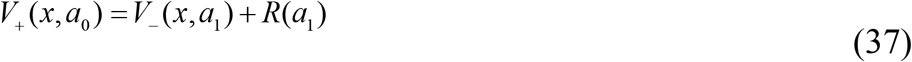

where *a*_0_ is a set of parameters, *a*_1_ is a function of *a*_0_, *R*(*a*_1_) is irrespective of *x*.

Following Gendenshtein [23] we call V_+_ and V_-_ are shape invariant. Evidently, the harmonic potential in present theory can be generalized to other potential that is shape-invariant with harmonic.

DNA replication can proceed only when the duplex is unwound. The speed of replication fork movement is 50000 bp/min for bacteria and about 2000 bp/min for eukaryotes [8]. That means the helix unwinding accompanying the replication should proceed in a very high rate. Topoisomerases play important role in helix unwinding. We conjecture that the microscopic mechanism of the topoisomerase may be related to the generation of a special kind of harmonic self-consistent field or its shape-invariant field.

### 6.3 Generalization to four kinds of bases

In a real DNA there are four kinds of bases,A,T,G and C. The symmetry of self-consistent field *V*(z) is strongly broken. The problem should be carefully studied. One important result is the annihilation and production operators 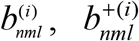 will be generalized to four kinds, namely 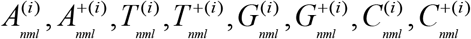 and bcorrespondingly the Hamiltonian Eq (15) should also be generalized. Note that there are two hydrogen bonds between A and T and three hydrogen bonds between C and G, the energy gap Δ in Eq (20) will be generalized to two, Δ_1_ and Δ_2_, namely

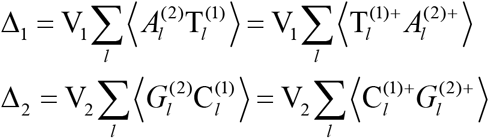

Accordingly, the quasi -particle can be generalized to four kinds.

### 6.4 Vertical interaction of base pairs and the sequence diversity

The W-C interaction between corresponding bases on two chains is called horizontal interaction. The stacking interaction between two neighboring bases on each chain is called vertical interaction. By use of annihilation and production operators the vertical interaction can be expressed as

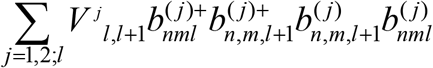

The vertical and horizontal interactions should be considered simultaneously. When the theory is generalized to four kinds of bases there are ten kinds of independent stacking interactions for DNA double chains. These stacking energies take quite different values, making one sequence differentiated from others. Moreover, the stacking energy is dependent of the local deviation of helix structure. Sometimes, one base mutation in a sequence could make a large variation of DNA structure. Therefore, the vertical interaction produces the sequence diversity and it makes possible that the sequence contains a large amount of biological information.

## Acknowledgement

The authors are indebted to Dr Yulai Bao and Ms Aiying Yang for their kind help in literature searching. The work was supported by Award Fund for Special Contributions to Science and Technology of Inner Mongolia Autonomous Region [No 2008 to LL].

## Appendix 1: Solution for 2D harmonic oscillator [14]

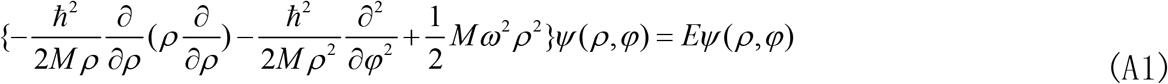

Set

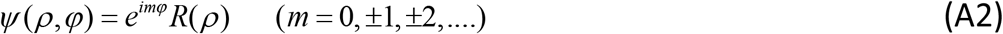

We obtain an equation for R(ρ).

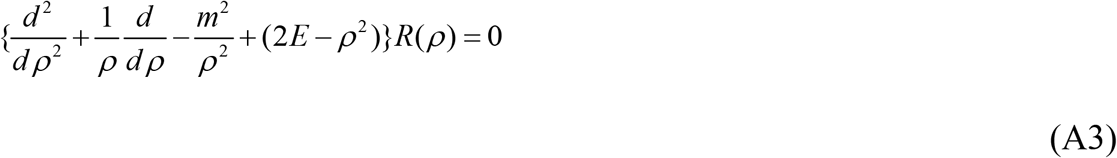

in natural unit of *ħ*=1, *m*=1 and ω=1.

By consideration of the boundary condition for ρ→0 and ρ→ ∞ we assume

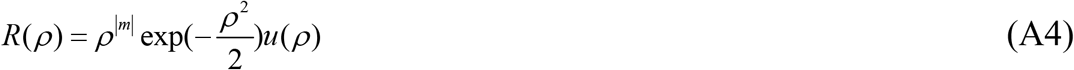

Inserting (A2)(A4) into (A1) we obtain

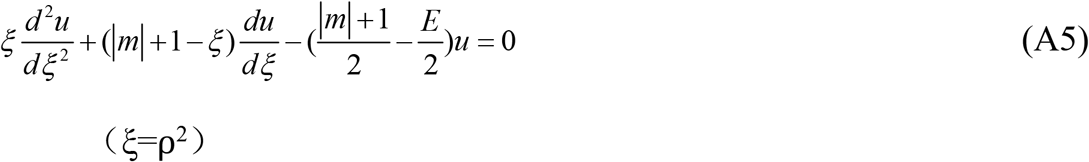

Eq(A5)is the confluent hyper-geometry equation exactly. To describe the bound state the eigenvalue E of Eq (A5) should take discrete values

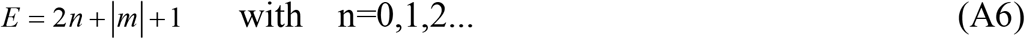

## Appendix 2: Calculation of the density of states

The quantum state in 2D harmonic potential is characterized by (n,m) and that in 1D periodic potential by Bloch wave number K or Wannier’s localized quantum number *l*. In tight binding model [17] [19]. we assume V_sym_(z) is a periodic well with width h - *b* and thick *b*

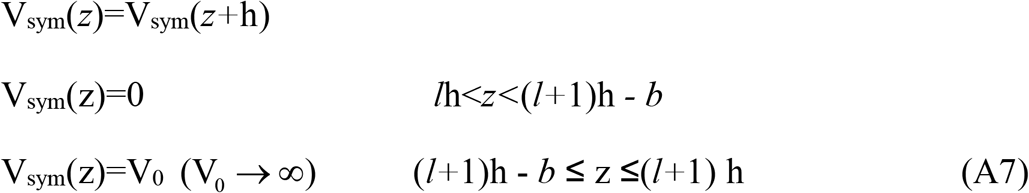

The normalized wave function of a particle with mass M in the *l*-th well

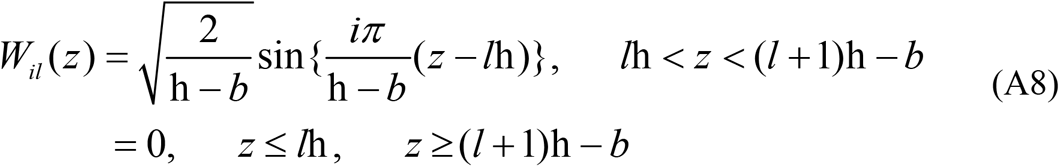

The energy level

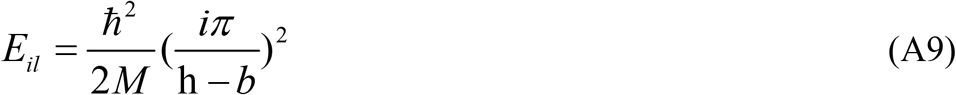

which is dependent of quantum number *i* (*i*=1, 2 …) only and degenerate with respect to *l*. The state density for small width *b*

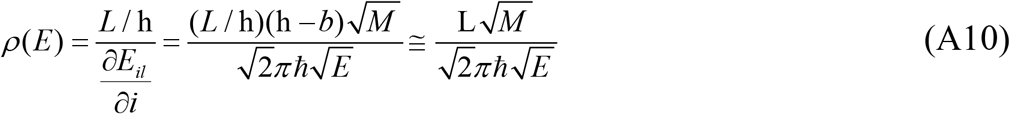

If the state density in energy band n=0, m=1 is considered then E in Eq (A10) should be replaced by ^*E* − 2*ħω*^, namely

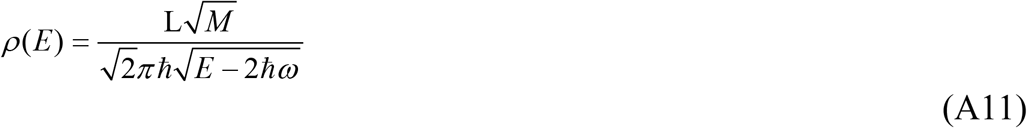

Note that the state density ((A10) and (A11)) is proportional to L but irrespective of the step of helix h.

## References

1. Wolynes, P.G. 2009. Some quantum weirdness in physiology. Proc Natl Acad Sci USA. 106, 17247–17248.

2. Lambert, N., Chen, Y.N., Cheng, Y.C., Li, C.M., Chen, G.Y. and Nori, F. 2013. Quantum biology. Nature Physics. 9, 10–18

3. Hiscock, H.G., Worster, S., Kattnig, D.R., Steers, C., Jin, Ye., Manolopoulos, D.E., Mouritsen H. and Hore, PJ. 2016. The quantum needle of the avian magnetic compass. Proc Natl Acad Sci USA. 113, 4634–4639.

4. Luo, L.F. 2014. Quantum theory on protein folding. Sci China Phys Mech & Astro. 57, 458–468.

5. Luo, L.F. and Lv, J. 2017. Quantum conformational transition in biological macromolecule. Quantitative Biology 5, 143–158. DOI 10.1007/s40484-016-0087-9.

6. Karplus M. 2014. Development of multiscale models for complex chemical systems from H+H2 to biomolecules (Nobel Lecture). Angewandte Chemie. 126 (38), 10152–10166.

7. Schrodinger E. 1944. What is life. Cambridge University Press.

8. Lewin B. 2008. Genes IX. Jones and Bartlett Pub. (Sudbury MA) Ch 31, Ch 18.

9. Chen X., Gu ZC., Wen XG. 2010. Local unitary transformation, long range quantum entanglement, wave function renormalization and topological order. Phys. Rev. B 82, 155138.

10. Liu CQ., Cao EH., Bai CL., Wang C., Liang SR. 2000. Polymorphism of Nucleic Acid Structure. High Education Press, China (Beijing) (in Chinese)

11. Heller DA. et al. 2006. Optical detection of DNA conformational polymorphism on single-walled carbon nanotubes. Science 311, 508–511.

12. Chakraborty A, Truhlar D G. 2005. Quantum mechanical reaction rate constants by vibrational configuration interaction: the OH + H2->H2O + H reaction as a function of temperature. Proc Natl Acad Sci USA 102, 6744–6749.

13. Yang GC. 2016. A quantum mechanical model of DNA. Talk at Int. Symposium on the Frontier of Big Data in Science. (Baotou)

14. Zeng JY. 2016. Quantum Mechanics (5^th^ Ed). Science Pub. (Beijing) (in Chinese).

15. Flugge S. 1974. Practical Quantum Mechanics. Springer-Verlag.

16. Panati G, Pisante A. 2013. Bloch bundles, Marzari-Vanderbilt functional and maximally localized Wannier functions. Communications in Math. Phys. 322(3), 835–875.

17. Huang K. 2014 Solid State Physics. Peking University Press.

18. Luo LF. 1987. Conformation dynamics of macromolecules. Int. J. Quantum Chemistry. 32, 435–450.

19. Economou EN. 2010. The Physics of Solids. Berlin-Heiderberg: Springer.

20. Huang KS. 1987. Statistical Mechanics (2^nd^ Edi). New York: Wiley.

21. Nelson P. 2004. Biological Physics: Energy, Information, Life. www.physics.upenn.edu/~pcn/

22. Nagapriya KS., Raychaudhuri AK. 2006. Direct observation of large temperature fluctuation during DNA thermal denaturation. Phys. Rev. Lett. 96, 038102.

23. Gendenshtein L. 1983 JETP Lett. 38, 356. (also discussed in [14]).

